# Longitudinal changes of ADHD symptoms in association with white matter microstructure: a tract-specific fixel-based analysis

**DOI:** 10.1101/2021.11.19.469248

**Authors:** Christienne G. Damatac, Sourena Soheili-Nezhad, Guilherme Blazquez Freches, Marcel P. Zwiers, Sanne de Bruijn, Seyma Ikde, Christel M. Portengen, Amy C. Abelmann, Janneke T. Dammers, Daan van Rooij, Sophie E. A. Akkermans, Jilly Naaijen, Barbara Franke, Jan K. Buitelaar, Christian F. Beckmann, Emma Sprooten

## Abstract

**Background:** Variation in the longitudinal course of childhood attention deficit/hyperactivity disorder (ADHD) coincides with neurodevelopmental maturation of brain structure and function. Prior work has attempted to determine how alterations in white matter (WM) relate to changes in symptom severity, but much of that work has been done in smaller cross-sectional samples using voxel-based analyses. Using standard diffusion-weighted imaging (DWI) methods, we previously showed WM alterations were associated with ADHD symptom remission over time in a longitudinal sample of probands, siblings, and unaffected individuals. Here, we extend this work by further assessing the nature of these changes in WM microstructure by including an additional follow-up measurement (aged 18 – 34 years), and using the more physiologically informative fixel-based analysis (FBA).

**Methods:** Data were obtained from 139 participants over 3 clinical and 2 follow-up DWI waves, and analyzed using FBA in regions-of-interest based on prior findings. We replicated previously reported significant models and extended them by adding another time-point, testing whether changes in combined ADHD and hyperactivity-impulsivity (HI) continuous symptom scores are associated with fixel metrics at follow-up.

**Results:** Clinical improvement in HI symptoms over time was associated with more fiber density at follow-up in the left corticospinal tract (lCST) (t_max_=1.092, standardized effect[SE]=0.044, *p*_FWE_=0.016). Improvement in combined ADHD symptoms over time was associated with more fiber cross-section at follow-up in the lCST (t_max_=3.775, SE=0.051, *p*_FWE_=0.019). **Conclusions**: Aberrant white matter development involves both lCST micro- and macrostructural alterations, and its path may be moderated by preceding symptom trajectory.

## 1. Introduction

Although (proto)typically considered a childhood syndrome, clinical trajectories of attention-deficit/hyperactivity disorder (ADHD) vary by individual. Many ADHD-affected adolescents exhibit improvement over time, but approximately two-thirds of them retain impairing symptoms into adulthood [1–3]. The neural substrates that determine this variable clinical course of childhood ADHD have been increasingly investigated through the years, yet the dynamic nature of these mechanisms in relation to maturation remains unclear. Theoretically, symptom remission occurs via brain compensation-reorganization, and/or normalization-convergence, with a possible fixed anomaly ‘scar’ or enduring neurological trait—all of which may concurrently arise in different brain regions [4]. In a double dissociative neurodevelopmental model of ADHD, the underlying neural mechanisms that control onset are distinct from those that drive remission [5]. Thus, onset can be characterized by dysfunctional subcortical structures remaining static throughout life, while remission may be separately associated with brain (particularly prefrontal cortex) maturation and compensation [4,6,7].

The theory that maturing frontal cortical regions compensate for initial childhood ADHD emergence via top-down regulatory processes, leading to eventual symptom remission, has been supported by magnetic resonance imaging (MRI) studies: Reduced symptom severity throughout development appears to correlate with prefrontal cortex maturation. White matter (WM) development in frontal-temporal areas subserving emotional and cognitive processes indeed continues to mature into early adulthood, coinciding with the typical age range of ADHD symptom remission [5,8–12]. Based on this model, it is possible to methodologically differentiate remitted from unaffected brains with MRI. Yet, previous neuroimaging studies have reported inconsistent results— perhaps because of study-specific differences (e.g. analysis methods, cross-sectional cohorts, sample characteristics). Considering this disorder’s neurodevelopmental component, sample age is especially important, making systematic longitudinal studies essential in deconstructing the etiological timeline of brain mechanisms in reference to remission.

Diffusion-weighted imaging (DWI) is an *in vivo* MRI method which measures the magnitude and direction of water molecules diffusing through brain tissue, reflecting the underlying architecture of axons and their ensheathing myelin. Diffusion tensor imaging (DTI) has been the most commonly used DWI method in ADHD studies, which have usually reported tensor-derived, voxel-wise measures like fractional anisotropy (FA). One follow-up case-control DTI investigation in men suggested that ADHD is a lasting neurobiological trait irrespective of remission or persistence: Compared to those who did not have childhood ADHD, probands with both remittent and persistent ADHD showed widespread reduced FA three decades post-diagnosis [6]. Others showed that children who exhibited symptom improvement had the most FA anomalies at follow-up [13]. However, given ADHD’s neurodevelopmental aspect, studying symptoms and brain tissues in late adolescence and early adulthood (as myelination continues) can give more relevant information about how remission is intertwined with maturation. While valuable, the few follow-up DTI reports to date were limited by categorical participant groups, sample characteristics (populations that were either pre-pubertal or well into adulthood), and only a single follow-up MRI measure—underscoring the need for more studies beyond the cross-sectional perspective.

A longitudinal design reveals temporal dynamics of underlying neurobiological processes and increases statistical power by reducing inter-subject variability. The NeuroIMAGE study and its latest follow-up, DELTA, is a longitudinal cohort of ADHD-affected probands, their siblings, and unaffected controls from childhood to adulthood [10,14–17]. We previously demonstrated that, at two different time-points and in two partly overlapping NeuroIMAGE samples, more improvement in combined ADHD and hyperactivity-impulsivity symptom scores over time were associated with lower FA at follow-up in an area where the left corticospinal tract (lCST) crosses the left superior longitudinal fasciculus (lSLF) [10,17]. In that same report, we also systematically demonstrated that symptom change is associated with neither baseline FA nor change in FA from baseline to follow-up. Symptom remission was counterintuitively and repeatedly associated with decreased FA later in life, whether from childhood to adolescence, or to early adulthood. Our longitudinal findings indicated divergent WM microstructure trajectories between individuals with persistent and remittent symptoms at a follow-up age range of 12 – 29 years. Now, we build on those previous results on the downstream effect of symptom progression on WM microstructure by asking whether the same relationship exists in the same brain areas at a later time window, when the cohort is aged 18 – 34 years (Figure 1).

**Figure 1.**
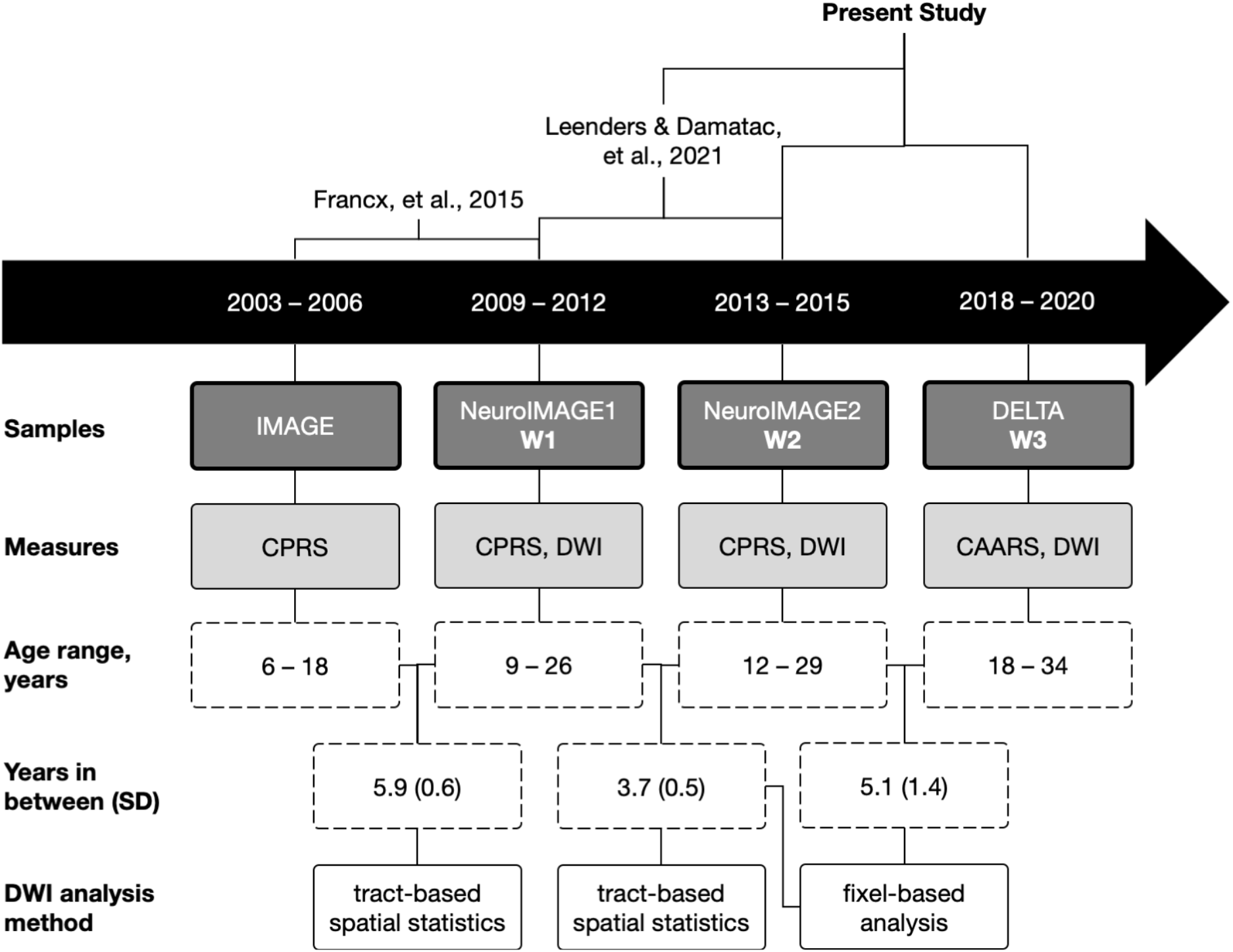
Schematic of how this study chronologically relates to previous studies, the samples included in each, relevant clinical and neuroimaging measurements, study sample age ranges, mean years (standard deviation) in between each acquisition wave, and the analysis methods used. The present study is a fixel-based analysis of W1 to W2 and W2 to W3, using only the models in which we found significant effects in a previous voxel-wise tract-based spatial statistical analysis of W1 to W2.

Although *in vivo* WM microstructure has been most commonly studied through tensor-derived metrics (e.g. FA), results from voxel-wise DTI-based methods can be unreliable or misleading in areas with complex fiber architecture [18]. Given that tensor-based reconstructions are an average across an entire voxel and that approximately 90% of voxels contain multiple fiber populations, we applied a high angular diffusion model: constrained spherical deconvolution [19]. Voxel-based methods that model crossing fibers (e.g. BEDPOSTX) only represent a subset of the full range of possible fiber orientation distributions (FODs), whereas constrained spherical deconvolution represents FODs as spherical harmonics, free to distinguish more or less arbitrary shapes [20]. Fixel-based analysis (FBA) applies the constrained spherical deconvolution model and can more accurately reconstruct a continuous FOD in both single- and multiple-fiber voxels—characterizing properties of each “fixel,” or specific fiber population in a voxel [21–25]. Fixels can be statistically analyzed for fiber-specific indices of underlying physiology: fiber density (FD), a microstructural measure of the within-voxel intra-axonal restricted compartment of a fiber population; fiber cross-section (FC), a macrostructural measure of the area perpendicular to the fiber orientation; and fiber density and cross-section (FDC), a combination of FD and FC [25]. Less FD can indicate axonal loss, while less FC can indicate macroscopic fiber atrophy [26–28]. FBA resolves crossing fibers more accurately as well as characterizes the microstructural and morphological, macrostructural properties of specific fiber populations.

One cross-sectional FBA showed that ADHD-affected children who had reduced fine motor competence also had lower WM microstructure in all three fixel metrics in the CST. These results suggest that cases had fewer and/or thinner CST axons, which may lead to reduced fiber bundle information transmission speed [29]. Despite the consistent clamor to resolve crossing fiber regions and FBA’s evident advantages, there have been no other published FBA applications in people with ADHD to our knowledge. Furthermore, besides our prior research, there have been no other longitudinal follow-up DWI studies of WM in the course of ADHD.

In an extension of our previous work in overlapping samples, here we followed 139 people over approximately 15 years. We used a more recent multi-shell DWI fiber model and a new follow-up measurement at an older age range. We aimed to assess the time-lag between the course of ADHD symptomatology and WM microstructure in *a priori* models and regions-of-interest. Because our smaller sample size at an older age range is not suitable for a data-driven search to discover any new relevant regions or tracts, the present analyses were intended to further understand the nature of our previous results. To compare FBA metrics to our previous FA findings, our first follow-up analysis used the same exact sample as our most recent longitudinal DTI study [17]. We hypothesized that, like in our earlier findings, ADHD symptom improvement would be associated with lower follow-up WM microstructure in the lSLF and lCST.

## 2. Methods

### 2.1 Participants

Clinical and MRI data were originally collected from probands with childhood ADHD, their first-degree relatives, and healthy families in one initial wave: NeuroIMAGE1 (W1) [14]. After an average of 3.7 years (standard deviation [SD] = 0.5 years), those participants were invited back for a second acquisition: NeuroIMAGE2 (W2). After a mean of 5.1 years (SD = 1.4 years), some individuals returned for another wave, DELTA (W3), which included only people who fulfilled full ADHD diagnostic criteria in at least one previous wave (Table 1). For the analyses here, we only included participants who had clinical data from at least two of the three waves and DWI data from W2 and/or W3 (Figure 1). For each time-point, there were no differences between the participants included in the current analyses and the complete sample in symptom severity, age, and sex (*p* > 0.12).

**Table 1.** Demographic and clinical characteristics of participants at Wave 1 (W1), Wave 2 (W2), and Wave 3 (W3) with mean and standard deviation (or numerical count and percentage). W3 included only those who fulfilled full ADHD diagnostic criteria in at least one previous wave. Values reported here are for all participants in the final sample after all quality control (N = 139). ^*a*^ IQ was estimated using the vocabulary and block design subtests of the Wechsler Intelligence Scale for Children or Wechsler Adult Intelligence Scale. ^*b*^ Medication ever used: Whether or not participants had ever taken ADHD medication. ^*c*^ Medications: Ritalin (methylphenidate), Concerta (methylphenidate), Strattera (atomoxetine), and any other ADHD medication. The majority of patients were taking prescription medication for ADHD, mostly methylphenidate or atomoxetine. ^*d*^ Symptom scores in W1 and W2 were collected via Conners’ Parent Rating Scale, and W3 scores were collected via the Conners’ Adult ADHD Rating Scale. ^*e*^ Combined symptom score: Sum of hyperactivity-impulsivity and inattention scores.

|  | Wave 1 |  | Wave 2 |  | Wave 3 |  | Test<br>Statistic | (p) |
| --- | --- | --- | --- | --- | --- | --- | --- | --- |
|  | N = 119 |  | N = 135 |  | N = 55 |  |  |  |
|  | Mean | (SD) | Mean | (SD) | Mean | (SD) |  |  |
| Age, years | 16.98 | (3.47) | 20.22 | (3.48) | 24.82 | (4.07) |  |  |
| Sex, female | N = 48 | 40% | N = 53 | 39% | N = 19 | 35% | $\chi^2_3 = 1.51$ | (0.68) |
| Estimated IQ<sup>a</sup> | 98.41 | (14.81) | 105 | (16.82) | 102.35 | (14.17) | $F_{2,192} = 2.23$ | (0.11) |
| Head motion,<br>framewise<br>displacement | 0.47 | (0.17) | 0.53 | (0.42) | 0.82 | (0.17) | $F_{2,298} = 24.71$ | (<10<sup>-10</sup>) |
| Handedness, right | N = 96 | 81% | N = 111 | 82% | N = 45 | 82% | $\chi^2_3 = 4.61$ | (0.20) |
| Medication ever<br>used, yes<sup>b,c</sup> | N = 66 | 56% | N = 45 | 33% | N = 42 | 76% | $\chi^2_3 = 32.03$ | (<10<sup>-6</sup>) |
| **Symptom raw**<br>**score by diagnostic**<br>**group<sup>d</sup>** |  |  |  |  |  |  |  |  |
| Combined score<sup>e</sup> | 13.03 | (12.33) | 12.96 | (13.05) | 20.53 | (9.34) |  |  |
| Hyperactivity-impulsivity score | 4.94 | (5.73) | 4.71 | (5.68) | 10.85 | (5.38) |  |  |
| Inattention score | 8.05 | (7.51) | 7.55 | (7.45) | 9.67 | (4.85) |  |  |
| **Comorbidity**<br>**diagnosis** |  |  |  |  |  |  |  |  |
| Anxiety disorder | N = 2 | 2% | N = 2 | 1% | N = 1 | 2% | $\chi^2_3 = 57.344$ | (<10<sup>-11</sup>) |
| Avoidant<br>personality disorder | N = 1 | 1% | N = 2 | 1% | N = 0 | 0% | $\chi^2_3 = 81.886$ | (<10<sup>-15</sup>) |
| Conduct disorder | N = 2 | 2% | N = 2 | 1% | NA | NA | $\chi^2_2 = 55.107$ | (<10<sup>-11</sup>) |
| Major depression | N = 1 | 1% | N = 1 | 1% | N = 4 | 7% | $\chi^2_3 = 57.435$ | (<10<sup>-11</sup>) |
| Oppositional<br>defiant disorder | N = 15 | 13% | N = 16 | 12% | NA | NA | $\chi^2_2 = 7.8901$ | (7.89) |
| Panic disorder | N = 0 | 0% | N = 1 | 1% | N = 0 | 0% | $\chi^2_3 = 137.76$ | (<10<sup>-15</sup>) |
| **Substance use** |  |  |  |  |  |  |  |  |
| Alcohol | N = 19 | 16% | N = 21 | 16% | N = 7 | 13% | $\chi^2_3 = 6.2621$ | (0.10) |
| Tobacco | N = 43 | 36% | N = 45 | 33% | N = 12 | 22% | $\chi^2_3 = 5.1952$ | (0.16) |
| Cannabis or hash | N = 22 | 18% | N = 23 | 17% | N = 6 | 11% | $\chi^2_3 = 6.8817$ | (0.08) |
| Other drugs | N = 6 | 5% | N = 6 | 4% | N = 2 | 4% | $\chi^2_3 = 23.734$ | (<10<sup>-4</sup>) |

Given our longitudinal design, we did not split our participants into cases versus controls. Through the years, symptom scores and diagnoses varied through time and participant characteristics changed from wave to wave (Figure 2). Some individuals originally recruited as controls or unaffected siblings developed ADHD at a later time point and others recruited as ADHD participants remitted, further highlighting the complex, variable course of ADHD. Alternative to a case-control categorization, ADHD can be operationalized as a continuous trait [30,31]. In a previous cross-sectional study, we systematically showed that, compared to categorical diagnoses, continuous symptom measures are more sensitive to diffusion-weighted brain features in this sample [16]. Thus, all models here used ADHD symptom scores, optimally capturing the dynamic and continuous nature of the ADHD spectrum throughout development in this longitudinal cohort.

**Figure 2.**
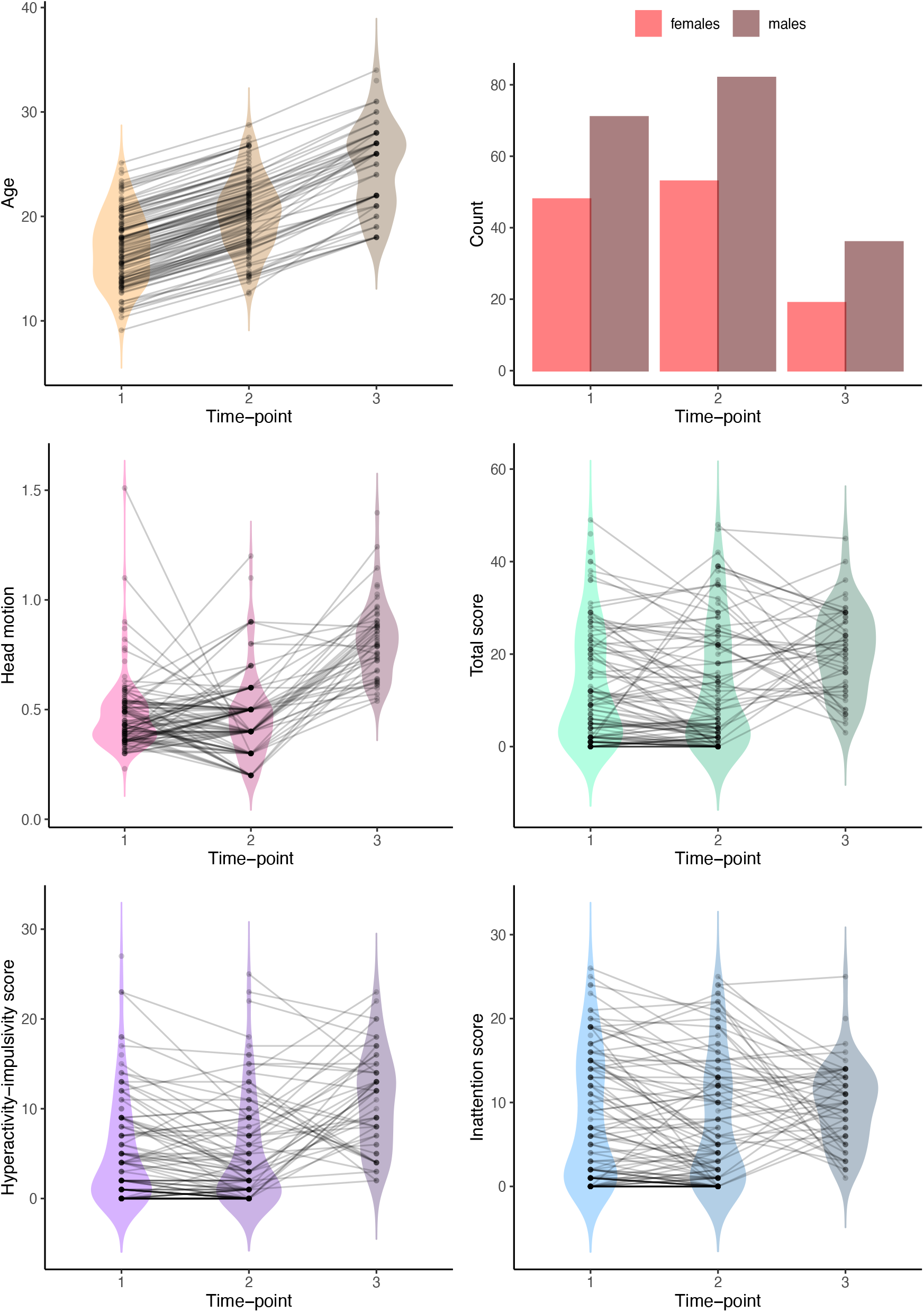
Change in participant characteristics from Wave 1, to Wave 2, to Wave 3. Longitudinal data points are connected by a line. Note that participants at W3 were selected on the basis of their history of ADHD diagnosis, so W3 tends to differ quite markedly from the other two waves, which also include never-affected controls. This conceals the typical pattern of average symptom remission that would be expected in a follow-up study without this selection criterion.

### 2.2 Clinical symptom measures

For continuous measures of symptom dimension severity and in accordance with our previous report, we used raw combined Conners’ Parent Rating Scale (CPRS) scores from W1 and W2, and Conners’ Adult ADHD Rating Scale (CAARS) scores from W3 for hyperactivity-impulsivity (HI) and inattention (IA) [17,32,33]. Here, we define symptom change (Δ) as the Conners’ score difference: Δscore = score_follow-up_ − score_baseline_

Baseline versus follow-up scores were always positively correlated with each other (Figure S1). A more positive Δ value indicates the worsening of symptoms, while a more negative Δ value indicates the improvement of symptoms over time. In this report, we refer to “symptom remission” dimensionally and not diagnostically, i.e. a decrease or improvement in symptom severity over time.

At W1 and W2, we assessed history of comorbid disorders with the Kiddie Schedule for Affective Disorder and Schizophrenia Present and Lifetime Version (K-SADS-PL) semi-structured interview [34,35]. For children aged <12 years, the child’s parents or the researchers assisted in completing the self-report questionnaires. Participants with elevated scores on ≥1 of the K-SADS-PL screening questions had to complete a full supplement for each disorder. At W3 (all participants were aged ≥18 years), we recorded history of comorbidity using the Structured Clinical Interview for DSM-5 Disorders (SCID-V) [36]. IQ was estimated using the vocabulary and block design subtests of the Wechsler Intelligence Scale for Children (WISC-III) or Wechsler Adult Intelligence Scale (WAIS-III). We excluded one whole dataset from a participant who had an estimated IQ <70. Our final sample’s demographic characteristics are summarized in Table 1.

### 2.3 Diffusion-weighted imaging acquisition, pre-processing, and quality control

At W2, single-shell DWI data were acquired with a 1.5-Tesla AVANTO scanner (Siemens, Erlangen, Germany) equipped with an 8-channel receive-only phased-array head coil using the following parameters: echo time/repetition time (TE/TR) = 97/8500 ms; GRAPPA-acceleration factor 2; voxel size = 2 × 2 × 2.2 mm; *b*-values = 0 (5 volumes, interleaved) and 1000 (60 directions) s/mm^2^; twice refocused pulsed-gradient spin-echo EPI; no partial Fourier. More details of this MRI data acquisition have been described previously [16,17]. Because our models only included follow-up neuroimaging data as a further investigation of the aforementioned analyses, we did not include W1 DWI.

At W3, multi-shell DWI data were acquired with a 3-Tesla Prisma scanner (Siemens, Erlangen, Germany) equipped with a 32-channel receive-only phased-array head coil using the following parameters: TE/TR = 75/2940 ms; multi-band acceleration factor = 3, voxel size = 1.8 mm^3^; *b*-values = 0 (11 volumes, interleaved), 1250 (86 directions), and 2500 (85 directions) s/mm^2^.

W2 and W3 images were pre-processed with MRtrix3 (version 3.0.1, http://www.mrtrix.org/) according to recommended FBA protocols for multi-shell data [24,25]. Pre-processing included denoising and unringing, motion and distortion correction, and bias field correction [37–43]. We visually inspected all corrected diffusion images and excluded whole datasets if any motion or distortion artefacts remained after pre-processing. After excluding 22 datasets, our final sample consisted of 154 total diffusion scans collected from 139 participants at W2 (N = 99) and W3 (N=55).

### 2.4 Fixel-based analysis

Following pre-processing, we computed two unique group average tissue response functions for W2: WM and cerebrospinal fluid (CSF) [44]. B0 images can be utilized like a second shell to estimate a CSF-specific response function for each participant [44]. By modeling distinct response functions for WM and CSF, we were able to enhance the signal from WM relative to CSF and include our single shell data in the multi-shell FBA pipeline. For W3, we calculate three response functions: WM, gray matter, and CSF [44]. We upsampled to 1.25 mm^3^ and performed multi-shell multi-tissue constrained spherical deconvolution on all images, resulting in a WM fiber orientation distribution (FOD) within each voxel [22,45]. Afterwards, we performed joint bias field correction and global intensity normalization for each of the multi-tissue compartment parameters [41]. We then separately generated two study-specific FOD population templates for W2 and W3 using 40 unrelated participants from each wave per template. Symptom scores did not differ between the individuals included in the population templates, versus those of the overall samples (P > 0.06).

For each population template, we calculated the FD, log(FC), and FDC (FDC = FD · FC) for each participant across all fixels. Instead of FC, we chose to calculate log(FC) so data would be centered around zero and normally distributed. The derivation of these fixel metrics, which are based on FOD lobe segmentation and subject-to-template registration warps, are described in detail elsewhere [46]. For each FOD template, we performed whole-brain fiber tractography and generated a fixel-fixel connectivity matrix from the whole-brain streamline tractograms.

To obtain each region-of-interest (left corticospinal tract: lCST; left superior longitudinal fasciculus: lSLFI, lSLFII, lSLFIII; right cingulum: rCG), we extracted the spherical harmonic peaks from each voxel of both FOD population templates. We then applied TractSeg, which is an automated convolutional neural network-based approach that directly segments tracts in fields of FOD peaks, circumventing any biases that may result from user-defined or atlas-based delineation [47]. Finally, we converted the resultant tractograms to fixel maps used as masks to constrain our search space during connectivity-based fixel enhancement [46] (Figure 3 and Figure S2).

**Figure 3.**
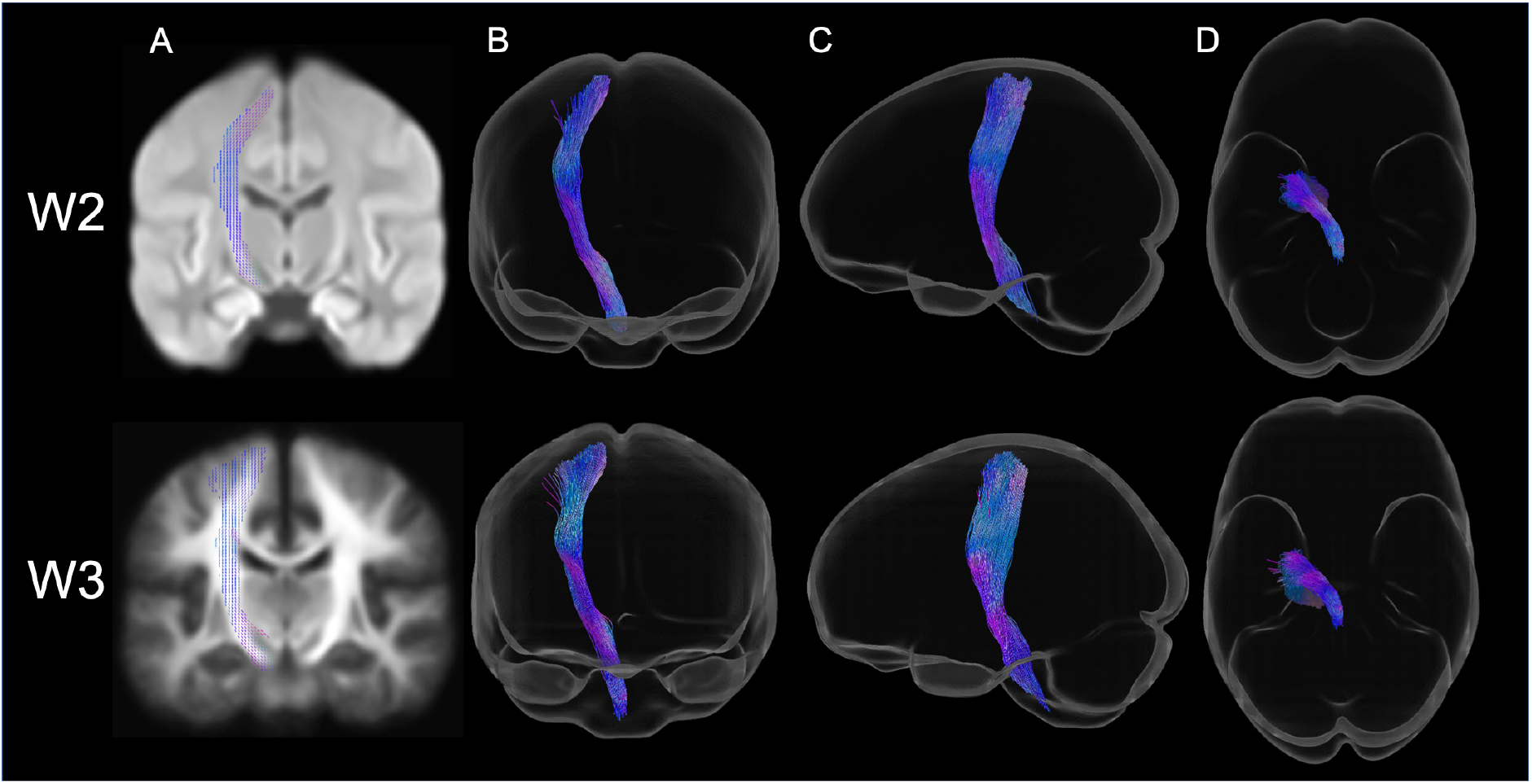
Tract-specific region-of-interest masks of the left corticospinal tract for Wave 2 (top) and Wave 3 (bottom) colored by direction (red: left-right, green: anterior-posterior, blue: inferior-superior). (A) Fixel mask overlaid on a single representative coronal slice of the study-specific white matter fiber orientation distribution template. Template contrast was adjusted and fixels have been thickened for visualization. (B) Coronal, (C) sagittal, and (D) axial views of the tract reconstruction from TractSeg (applied to each fiber orientation distribution template) and displayed in glass brains for visualization.

### 2.5 Statistical analyses

To control for the lack of independence in our sample due to siblings, we designed multi-level exchangeability blocks per wave and used FSL PALM to generate a set of 5000 permutations per wave [48,49]. Our blocks did not allow permutation between all individuals; instead, we constrained permutations at both the whole-block level (i.e. permute between families of the same size) and within-block level (i.e. permute within families) (Figure S3). We used each set as an input for its respective wave to define permutations in data shuffling during nonparametric testing.

We demeaned our design matrices using *Jmisc* in *R* (version 4.0.2) and applied connectivity-based fixel enhancement to the fixel-fixel connectivity matrices using smoothed fixel data [46]. Using only models in which we previously found significant effects (i.e. not IA, but only HI and combined scores), for each fixel metric and each tract region-of-interest, we constructed general linear models (GLMs) to separately test whether combined ADHD or HI symptom score change (Δscore as independent variables) are associated with fixel metrics at follow-up (as dependent variables) [17]. Our covariates were: symptom score (either combined or HI) at baseline, change in age (Δage = age _follow-up_ − age _baseline_), age at baseline, sex, and head motion (framewise displacement) at follow-up. For the W2 FBA, follow-up was W2 and baseline was W1, while for the W3 FBA, follow-up was W3 and baseline was W2: fixel metric _follow-up_ ~ Δscore + score _baseline_ + Δage + age _baseline_ + sex + head motion _follow-up_

As a secondary cross-sectional analysis in W3 only, using the same aforementioned methods, we tested for an effect of CAARS score (IA, HI, and combined) on each of the three fixel metrics at the level of the whole brain, as well as rCG only. We used age, sex, and head motion (framewise displacement) as covariates: fixel metric ~ score + age + sex + head motion

Fixels were considered statistically significant at family-wise error-corrected *p* less than 0.05 (*p*_FWE_ < 0.05).

## 3. Results

### 3.1 Association between white matter at Wave 2 and the change in symptoms from Wave 1 to Wave 2

W2 fiber density (FD) in the lCST was significantly negatively associated with ΔHI score (*t*_max_ = 1.092, standardized effect [SE] = 0.044, *p*_FWE_ = 0.016; Figure 4). There were no other significant associations between WM microstructure in any other tracts at follow-up and Δcombined or ΔHI score (all *p*_FWE_ ≥ 0.051; Table S1).

**Figure 4.**
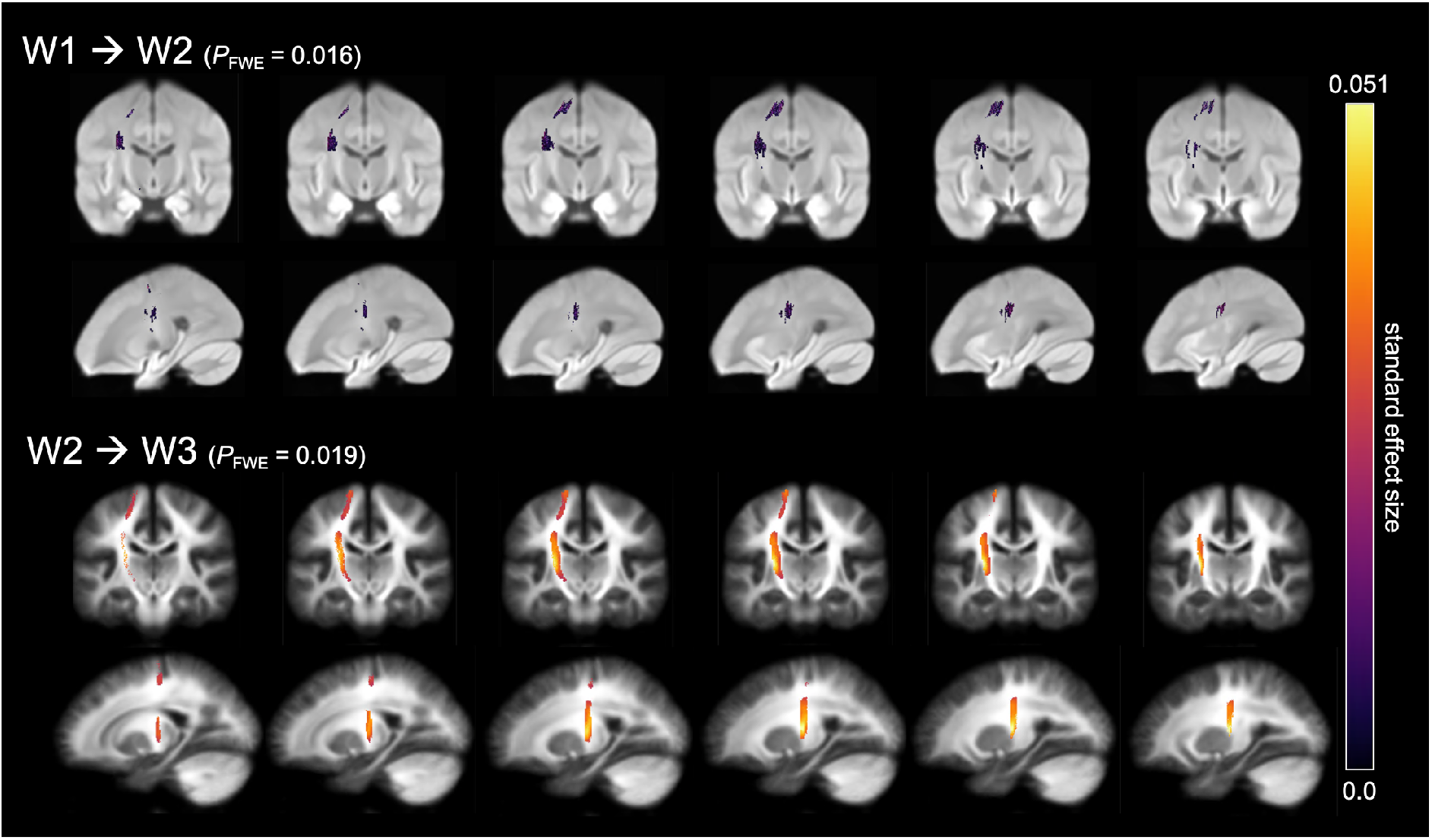
Symptom change may precede lCST WM microstructure plasticity. **Top**: Improvement of HI score is associated with more follow-up fiber density (FD). **Bottom**: Improvement of combined score is associated with more follow-up fiber cross-section (FC). Streamline segments have been cropped from the template tractogram to include only streamline points that correspond to significant fixels for this tract (FWE-corrected *p*-value < 0.05). Significant streamlines are colored by the standard effect size of ‘Δscore’ on ‘FD at W2’ and ‘log(FC) at W3’ and displayed across coronal and sagittal slices of the study-specific white matter fiber orientation distribution templates.

### 3.2 Association between white matter at Wave 3 and the change in symptoms from Wave 2 to Wave 3

W3 log of fiber cross-section (log[FC]) in the lCST was significantly negatively associated with Δcombined symptom score (*t*_max_ = 3.775, SE = 0.051, *p*_FWE_ = 0.019; Figure 4). There were no other significant associations between WM microstructure at follow-up in lSLFI, lSLFII, and lSLFIII and Δcombined or ΔHI symptom score (all *p*_FWE_ ≥ 0.058; Table S2).

### 3.3 Association between white matter and symptoms at Wave 3 only

None of the global or rCG WM microstructure metrics were significantly associated with HI, IA, or combined CAARS score cross-sectionally (all *p*_FWE_ ≥ 0.053; Table S3).

## 4. Discussion

We conducted a unique study of WM microstructure and longitudinal ADHD symptom development between ages 9 and 34 years. Using the FBA framework, we discovered two findings in the lCST: (1) HI symptom improvement was associated with axonal expansion at follow-up, and (2) combined ADHD symptom improvement was associated with a larger total cross-sectional area at follow-up at a slightly later age-range. Initially, a previous voxel-wise analysis in an overlapping sample found that improved HI symptoms were associated with lower follow-up FA (W1, aged 9 – 26 years) [10]. Subsequently, we extended this sample by adding a second DWI time-point (W2, aged 12 – 29 years), and systematically applied and excluded specific models—ultimately replicating the same effects on follow-up FA in the same WM region [17]. Given the counterintuitive nature of these previous highly consistent results, the present analysis aimed to further understand the physiological origins and its dynamic nature in relation to maturation. Thus here, in the exact same sample (W1-W2) and including yet a third DWI acquisition (W3, aged 18 – 34 years), using the more advanced FBA method, and employing the same GLMs in which we previously found significant voxel-wise effects, we have found increased FD in relation to HI remission, and increased FC in relation to combined symptom remission, in only the lCST and not the lSLF. In contrast to our previous finding using DTI-based methods, our current finding using FBA is more intuitive in the direction of its effect: Indices that are generally indicative of “stronger” fibers were associated with clinical improvements over time.

The fixel metrics we used for quantifying WM microstructure contain complementary information. FD is thought to be related to the microstructural properties of WM, whereas FC pertains to the macrostructural properties (cross-sectional area). FD is an estimate of the intracellular volume of fibers oriented in a particular direction. Higher FD at follow-up could result from developmental processes like axon diameter growth, or more axons occupying a given space [50]. In our W1-to-W2 analysis, greater lCST axonal density in individuals who became less hyperactive-impulsive over time suggests plasticity, or a greater ability to relay information, after symptom improvement. Furthermore, higher myelin content could decrease the water exchange between intra- and extra-axonal compartments, resulting in an apparent increase in the intra-axonal compartment’s volume and, hence, an increase in FD [42,51]. FC measures the morphological macroscopic change in the cross-sectional area perpendicular to a fiber bundle (calculated during registration to the template image). In W2-to-W3, higher follow-up lCST cross-sectional area in individuals whose combined symptom score improved, again, suggests plasticity, greater myelination, or fiber bundle organization after symptom remission [25].

Although the direction of effects in our FBA analyses are opposite to that of our aforementioned FA analyses, they are not incompatible. In some cases, crossing fiber complexity can have an inverse correlation with FA, wherein greater complexity occurs when more fixels in a voxel have the same fiber density [52]. An analogous inverse association exists in our previous W1-to-W2 voxel-wise analysis, wherein less follow-up FA was associated with improved HI symptom score. Notably, our results were in the approximate location of where the lSLF and lCST cross, while our present fixel-wise results in the lCST seem to be absent from where the lSLF crosses this tract. Therefore, as we previously suggested, our tract-based spatial statistics results may have been due to the neuroanatomical location of the effects, which, when labeled with an atlas, were in an area where these tracts cross. Compared to our voxel-wise study, we presently accounted for crossing fibers better through FBA, as well as the specific, FOD-based segmentation of these tracts as separate regions-of-interest. Accordingly, symptom improvement over time can conceivably be associated with increased CST fiber maturation, which by our previous DTI methods may have appeared as reduced FA in voxels where a more dominant SLF crosses those corticospinal fibers.

The lCST is the only tract in which we have consistently found longitudinal effects. However, a cross-sectional study of symptoms in an overlapping W1 sample found the most FA differences in the right cingulum-angular bundle [16]. Anatomically, this differs from that of the present and previous longitudinal effects, which suggests a dissociation in the WM tracts associated with cross-sectional differences versus those that are associated with symptom remission. This dissociation points to the neurodevelopmental models of remission, which all predict atypical neural features in adults with persistent ADHD, but have different predictions about those with remittent symptoms: If symptom remission occurred via WM normalization, convergence, or passive delayed maturation, then we would have observed neurological alterations at follow-up in participants with persistent symptoms, but no differences between those with remittent symptoms and healthy controls. If, regardless of symptom trajectory, this disorder imparted an indelible mark or scar on the brain, then we would not have observed follow-up neurological differences between those with persistent versus remittent symptoms, and only healthy controls would have been differentiated by our follow-up analyses. If symptom remission occurred via compensation or reorganization, then remitted brains would have differed from both the never affected and the persistent ADHD brains, but in different ways. The dissociation we observed in the tract that is important for symptom remission versus the tract that is important for symptom severity implies the last model of remission. According to our current findings, we tentatively suggest an interpretation consistent with our previous report: In a top-down fashion, remitters may have learned compensatory strategies to overcome symptoms as they aged, while persisters may have either learned disadvantageous strategies, other beneficial (but insufficiently effective) compensatory strategies, or none at all—leading to diverging WM development trajectories in specific brain regions in individuals with persistent ADHD symptoms (Figure S4).

Based on our longitudinal design, we postulate that different WM alteration patterns are associated with symptom trajectory in a tract-specific manner: HI symptom remission preceded lCST plasticity at 20 years median age (range 12 – 29 years), and combined symptom remission preceded lCST plasticity at 26 years median age (range 18 – 34 years). Perhaps our sample at a slightly younger age, in response to HI symptom improvement or learning new skills, gained more lCST fibers over time. Tract expansion could have been a compensatory mechanism to improve motor control, followed by more myelination of those fibers. Then, as our participants became slightly older, improvement in both dimensions may have led to greater lCST WM microstructure and improved motor control (Figure S5). We can speculate that improved IA (and related executive control) could help suppress HI, leading to greater motor control evinced as larger FC at a later age. In our remitters, higher measures of lCST WM might also result from reorganization in other brain areas outside of the tracts we studied. Along with our previous study, we have again found that alterations in WM microstructure appear to follow symptom improvement. Speculatively, this suggests that WM changes may be a downstream result of ADHD symptom remission.

A strength of the current study is its large sample size over three clinical and two DWI time-points. Our approach using two separate follow-up analyses lent further characterization to the temporal dynamics of ADHD-WM microstructure interplay. Of particular concern given this disorder, we mitigated potential confounding effects of head motion through careful data screening, correction during preprocessing, and inclusion as a covariate in our models. Using multi-shell FBA, we demonstrated that WM-associated differences are fiber-specific even within regions of crossing fibers, and we were able to further characterize WM micro- and macrostructural properties. Nonetheless, a limitation of this study is that the first follow-up analysis included DWI data acquired with only one relatively low b-value at 1.5T, which may have precluded us from discovering effects in other tracts and/or symptom dimensions since higher b-value shells improve correspondence between FD estimates and intra-axonal signal fraction simulations by increasing extra-axonal signal suppression [53]. Second, our inability to replicate our previous cross-sectional tractography findings may stem from the different DWI analysis methods used; also, W3 included a comparatively smaller number of affected-only individuals, which reduced our statistical power to detect smaller cross-sectional effects in the cingulum. Third, our follow-up samples were prone to selection bias from attrition and our explicit selection criteria in W3, where we also used different instruments (CRPS vs. CAARS) and raters. Returning participants were different from those who participated only once. Finally, even in a longitudinal study, we cannot prove causality. ADHD symptom persistence is likely associated with many other factors in daily life, or medication, or comorbid symptomatology—and any combination of these could also contribute to neurological differences at follow-up. Given our small sample size, we used *a priori* regions-of-interest and models, but a larger whole-brain analysis would be a more sufficient test of which mechanisms of remission are at play.

Our findings contribute to the growing body of evidence describing the progression of symptoms in relation to WM development. Defining the correlates and predictors of remission may eventually lead to an improved allocation of treatment resources for persistent or complicated ADHD. A better understanding of the underlying neural mechanisms of these changes in time can contribute to the promotion of favorable future perspectives for children and adolescents with this disorder.

## Abbreviations

ADHD: attention-deficit hyperactivity disorder
DTI: diffusion tensor imaging
DWI: diffusion-weighted imaging
FA: fractional anisotropy
FBA: fixel-based analysis
FC: fiber cross-section
FD: fiber density
FDC: fiber density and cross-section
FOD: fiber orientation distribution
HI: hyperactivity-impulsivity
IA: inattention
lCST: left corticospinal tract
lSLF: left superior longitudinal fasciculus
MRI: magnetic resonance imaging
SE: standardized effect
WM: white matter

## 5. Funding, acknowledgments, and financial disclosures

The authors would like to thank all of the families who participated in this study and all of the researchers who collected the data. This study sample is from the NeuroIMAGE project. NeuroIMAGE is the longitudinal follow-up study of the Dutch part of the International Multisite ADHD Genetics (IMAGE) project, which was a multi-site, international effort. NeuroIMAGE was supported by an NWO Large Investment Grant 1750102007010 and NWO Brain & Cognition an Integrative Approach Grant (433-09-242) (to J.K.B.), and grants from Radboud University Nijmegen Medical Center, University Medical Center Groningen and Accare, and VU University Amsterdam. Funding agencies had no role in study design, data collection, interpretation or influence on writing. J.N. is supported by an NWO Veni grant (no. VI.Veni.194.032). B.F. has received educational speaking fees from Medice. J.K.B. has been in the past 3 years a consultant to / member of advisory board of / and/or speaker for Takeda/Shire, Roche, Medice, Angelini, Janssen, and Servier. He is not an employee of any of these companies, and not a stock shareholder of any of these companies. He has no other financial or material support, including expert testimony, patents, royalties. All other authors report no biomedical financial interests or potential conflicts of interest.

## 8. Supplementary material

**Table S1.** Results from the tract-specific fixel-based analysis of W1 to W2 in lCST and lSLF.

| Search space | Dependent variable<br>Fixel-based metric | Contrast | Independent variable<br>Symptom score change<br>($\Delta = W2 - W1$) | $t_{\max}$ | Std. effect | $P_{FWE}$ |
| --- | --- | --- | --- | --- | --- | --- |
| Left corticospinal tract | Fiber density | ( + ) | combined | 0.858 | 0.011 | 0.165 |
|  |  | ( - ) | combined | 0.917 | 0.012 | 0.104 |
|  |  | ( + ) | HI | 0.949 | 0.038 | 0.225 |
|  |  | ( - ) | HI | **1.092** | **0.044** | **0.016\*** |
|  | Fiber cross-section | ( + ) | combined | 1.846 | 0.023 | 0.591 |
|  |  | ( - ) | combined | 2.264 | 0.029 | 0.343 |
|  |  | ( + ) | HI | 1.909 | 0.076 | 0.627 |
|  |  | ( - ) | HI | 2.189 | 0.087 | 0.536 |
|  | Fiber density and cross-section | ( + ) | combined | 0.969 | 0.012 | 0.280 |
|  |  | ( - ) | combined | 0.828 | 0.011 | 0.439 |
|  |  | ( + ) | HI | 1.051 | 0.042 | 0.294 |
|  |  | ( - ) | HI | 0.948 | 0.038 | 0.083 |
| Left superior longitudinal fasciculus I | Fiber density | ( + ) | combined | 2.238 | 0.028 | 0.496 |
|  |  | ( - ) | combined | 1.300 | 0.017 | 0.285 |
|  | Fiber cross-section | ( + ) | combined | 2.627 | 0.033 | 0.252 |
|  |  | ( - ) | combined | 1.759 | 0.023 | 0.474 |
|  | Fiber density and cross-section | ( + ) | combined | 2.483 | 0.032 | 0.112 |
|  |  | ( - ) | combined | 1.351 | 0.017 | 0.493 |
| Left superior longitudinal fasciculus II | Fiber density | ( + ) | combined | 1.547 | 0.020 | 0.462 |
|  |  | ( - ) | combined | 1.462 | 0.019 | 0.682 |
|  | Fiber cross-section | ( + ) | combined | 1.987 | 0.025 | 0.319 |
|  |  | ( - ) | combined | 1.372 | 0.017 | 0.625 |
|  | Fiber density and cross-section | ( + ) | combined | 1.666 | 0.021 | 0.245 |
|  |  | ( - ) | combined | 1.304 | 0.017 | 0.777 |
| Left superior longitudinal fasciculus III | Fiber density | ( + ) | combined | 1.464 | 0.019 | 0.762 |
|  |  | ( - ) | combined | 1.290 | 0.016 | 0.225 |
|  | Fiber cross-section | ( + ) | combined | 1.331 | 0.017 | 0.709 |
|  |  | ( - ) | combined | 3.121 | 0.040 | 0.051 |
|  | Fiber density and cross-section | ( + ) | combined | 1.766 | 0.022 | 0.863 |
|  |  | ( - ) | combined | 1.239 | 0.016 | 0.407 |

**Table S2.**
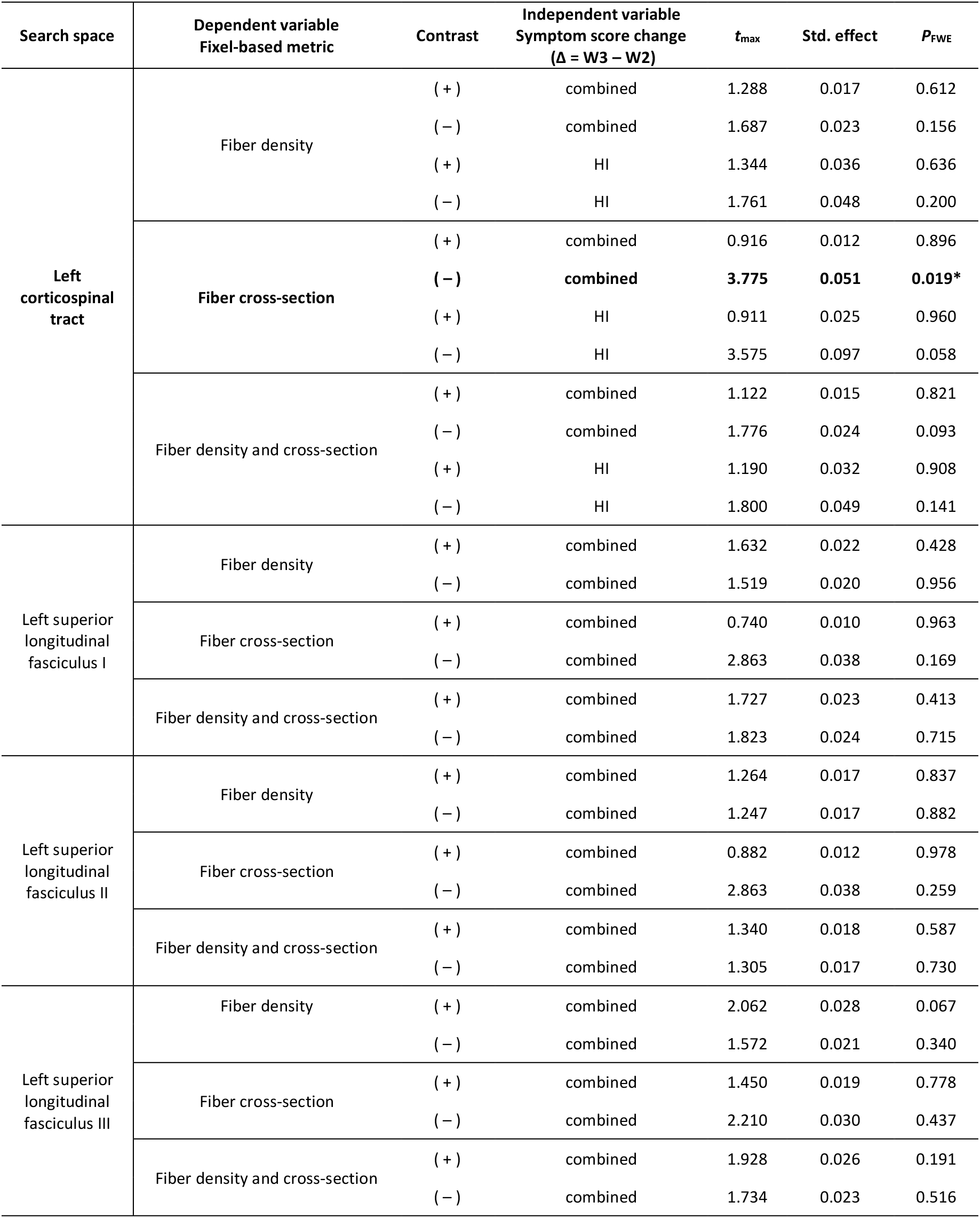
Results from the tract-specific fixel-based analysis of W2 to W3 in lCST and lSLF.

**Table S3.**
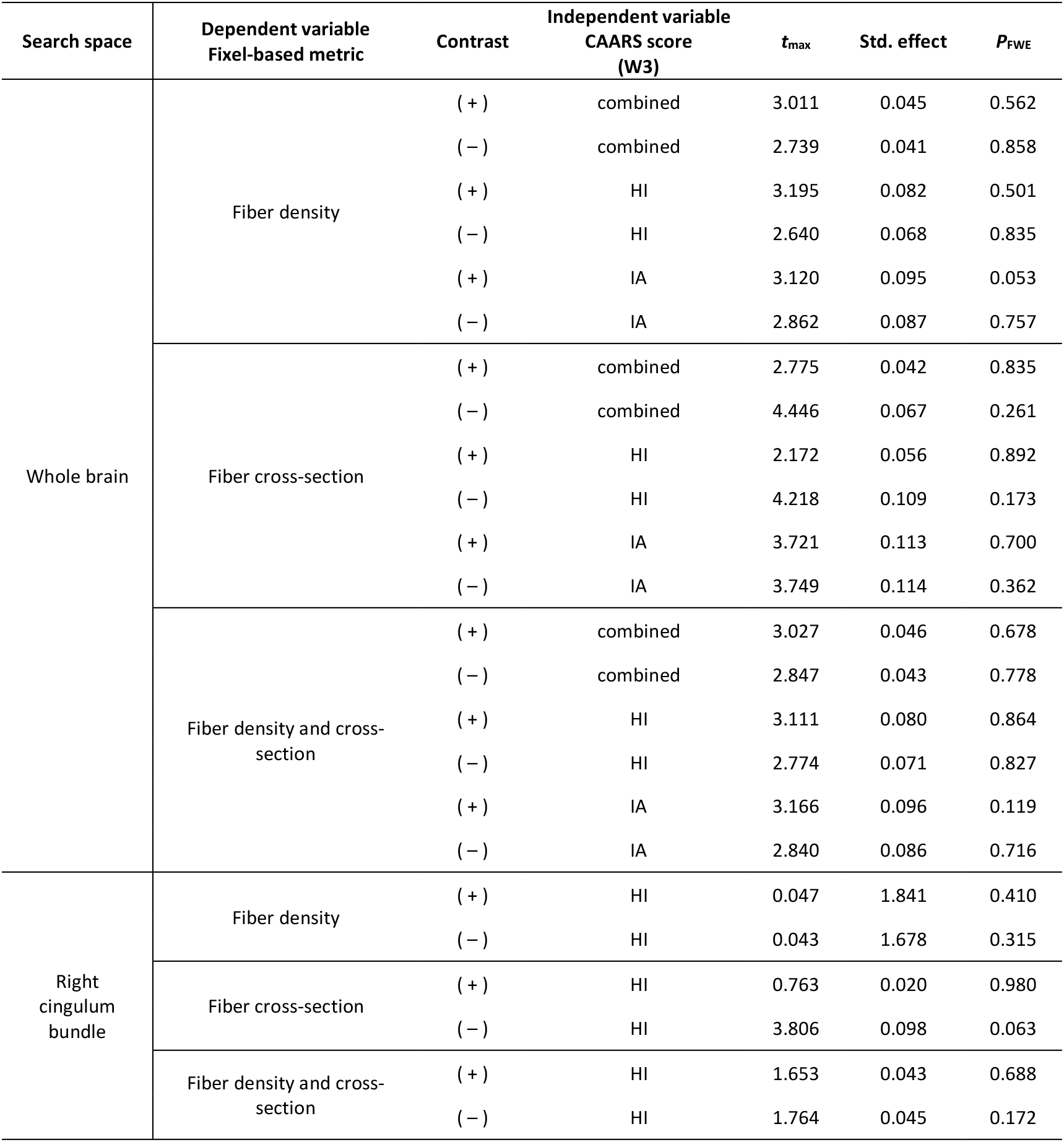
Results from the cross-sectional global and tract-specific fixel-based analysis of W3 only. All general linear models showed no significant effects of CAARS score on any fixel-based metric.

**Figure S1.**
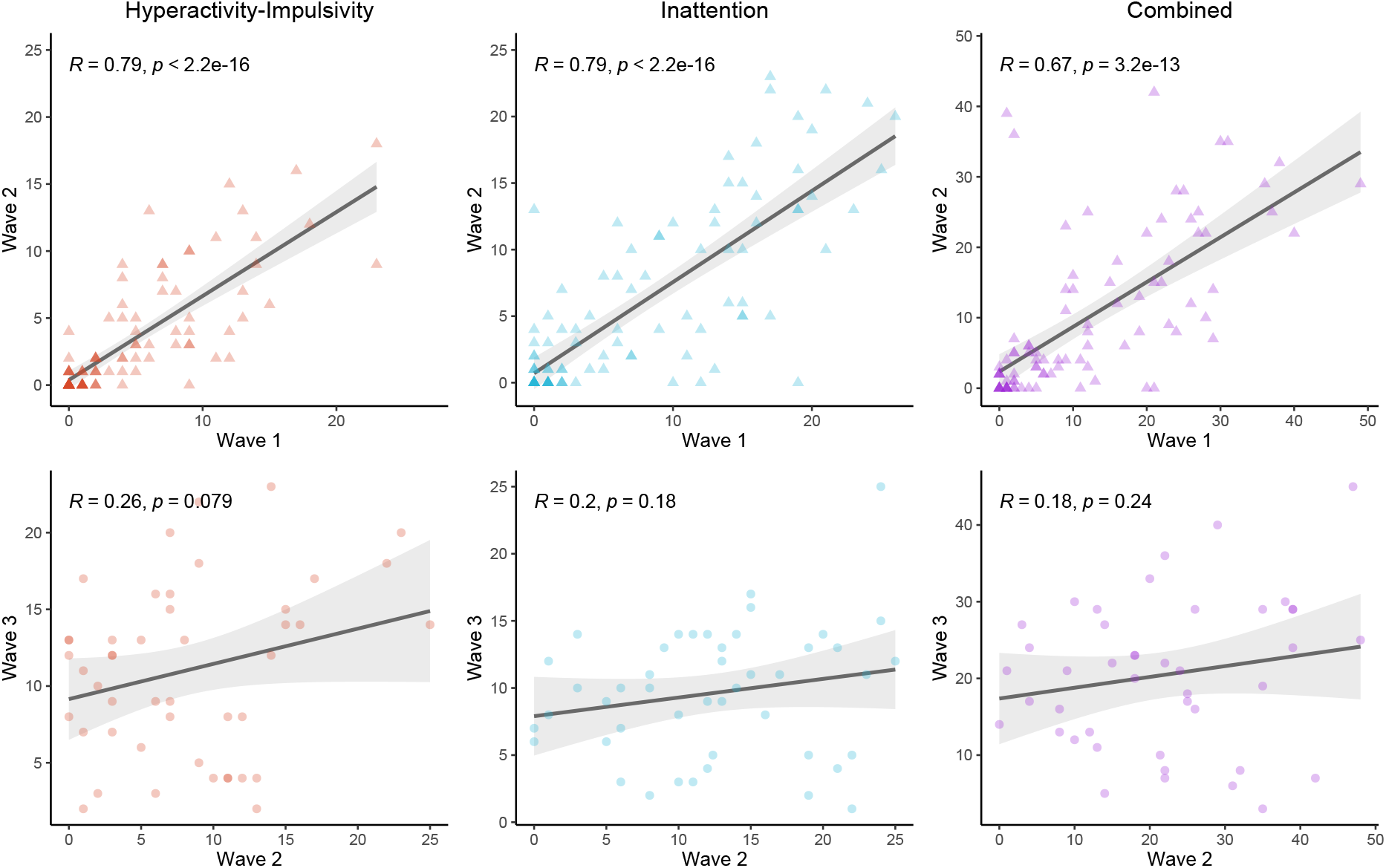
Correlation scatterplots of baseline against follow-up scores colored by dimension with 95% confidence intervals and Pearson correlation coefficients reported for each. **Top row**: Wave 1 associations with Wave 2 scores. **Bottom row**: Wave 2 associations with Wave 3 scores. **Left column**: Hyperactivity-impulsivity dimension scores. **Center column:** Inattention dimension scores. **Right column**: Combined scores (calculated as the sum of hyperactivity-impulsivity and inattention). Darker colored points indicate individuals with overlapping score data.

**Figure S2.**
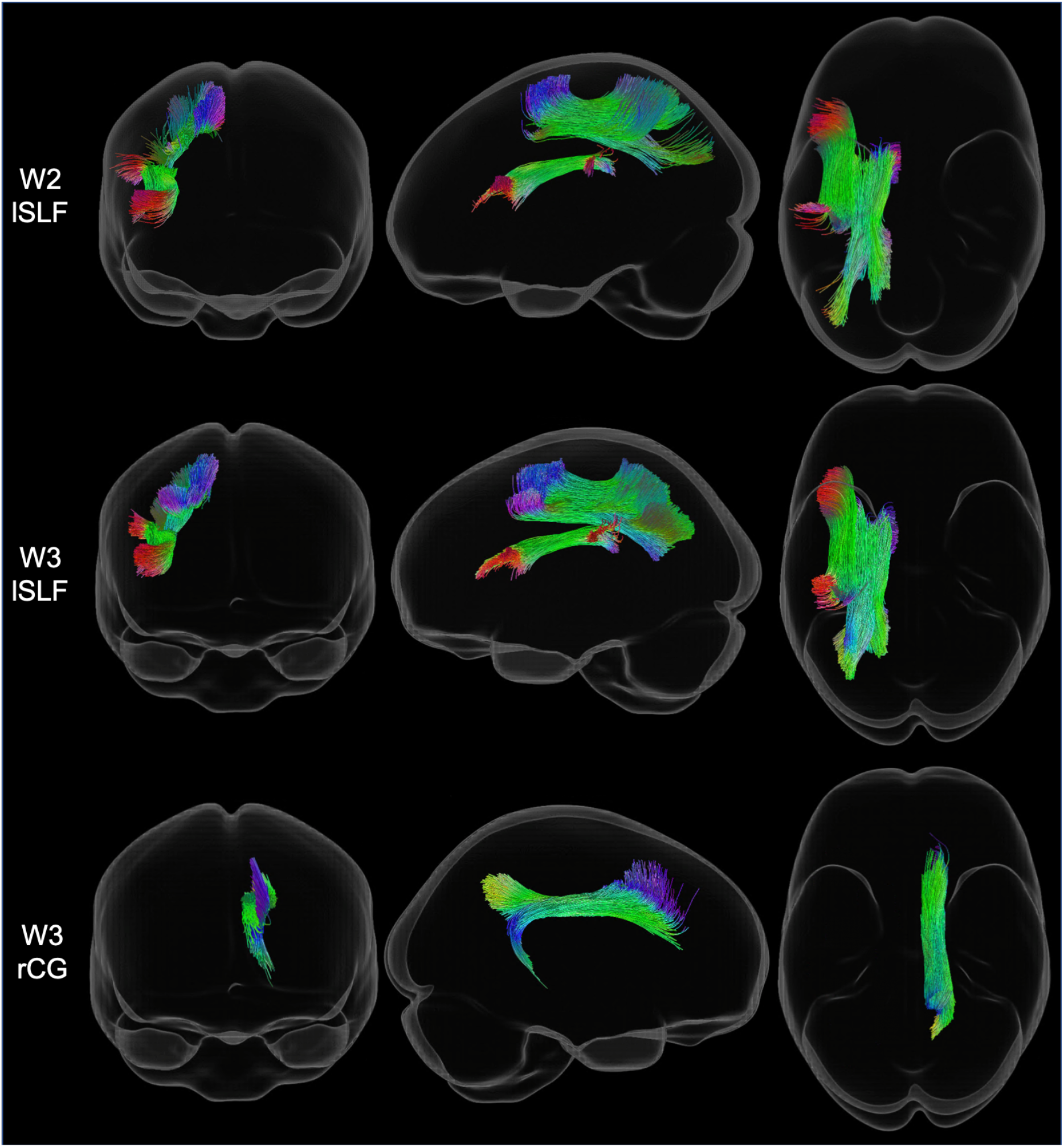
Tract-specific region-of-interest masks of the left superior longitudinal fasciculus (lSLF) at Wave 2 and Wave 3, and the right cingulum bundle (rCG) at Wave 3 colored by direction (red: left-right, green: anterior-posterior, blue: inferior-superior). Coronal (left column), sagittal (middle column), and axial (right column) views of the tract reconstructions from TractSeg (applied to the fiber orientation distribution templates) and displayed in glass brains for visualization.

**Figure S3.**
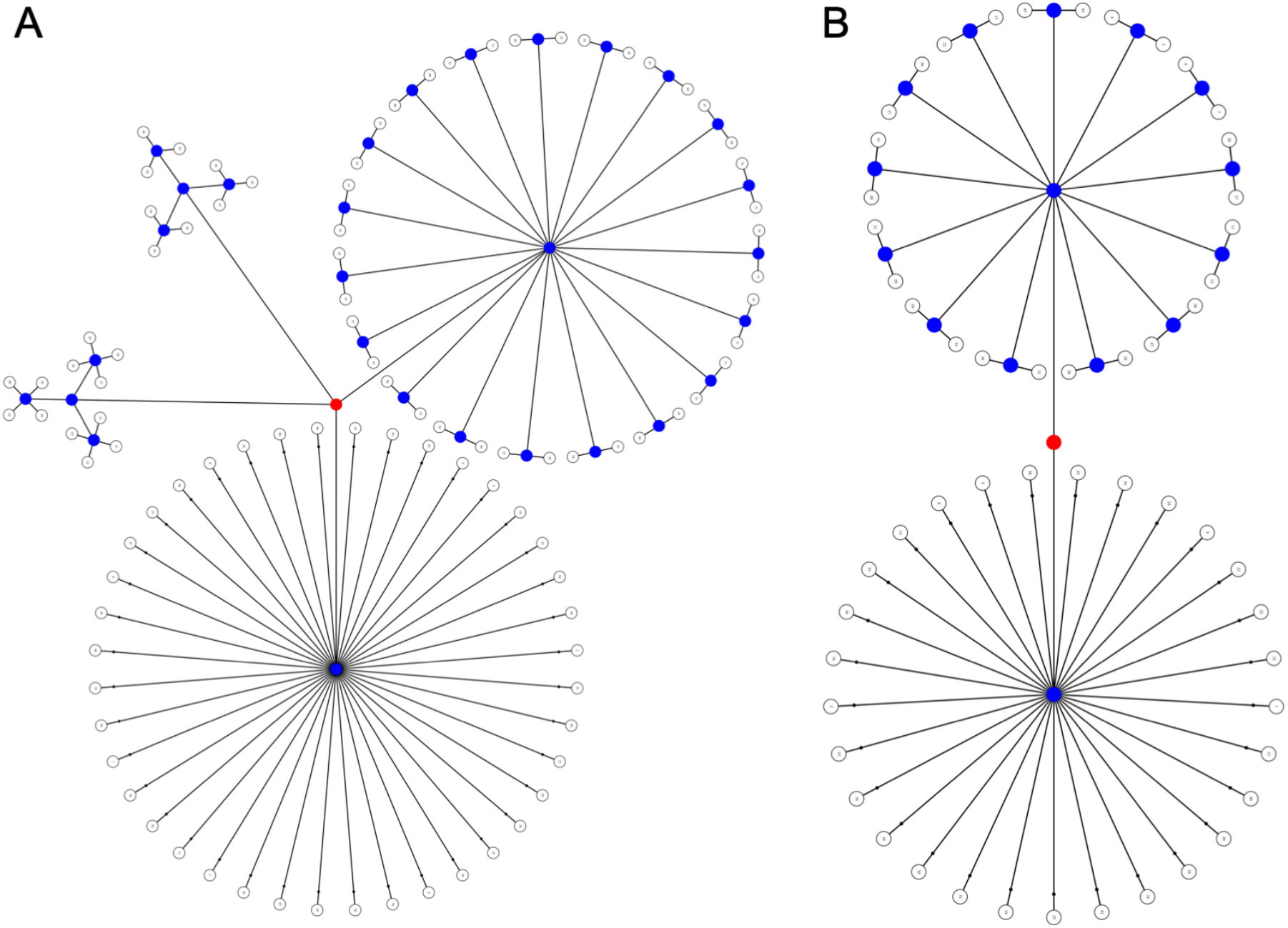
Visualization of the exchangeability block structure of (A) Wave 2 and (B) Wave 3, each represented in permutation trees. Each white dot is one individual and each group of white dots represents families of a specific size. At each permutation, branches beginning at blue dots can be permuted, while those beginning in red dots cannot. These permutation sets were generated with FSL PALM and used in connectivity-based fixel enhancement analysis to control for related siblings in each sample.

**Figure S4.**
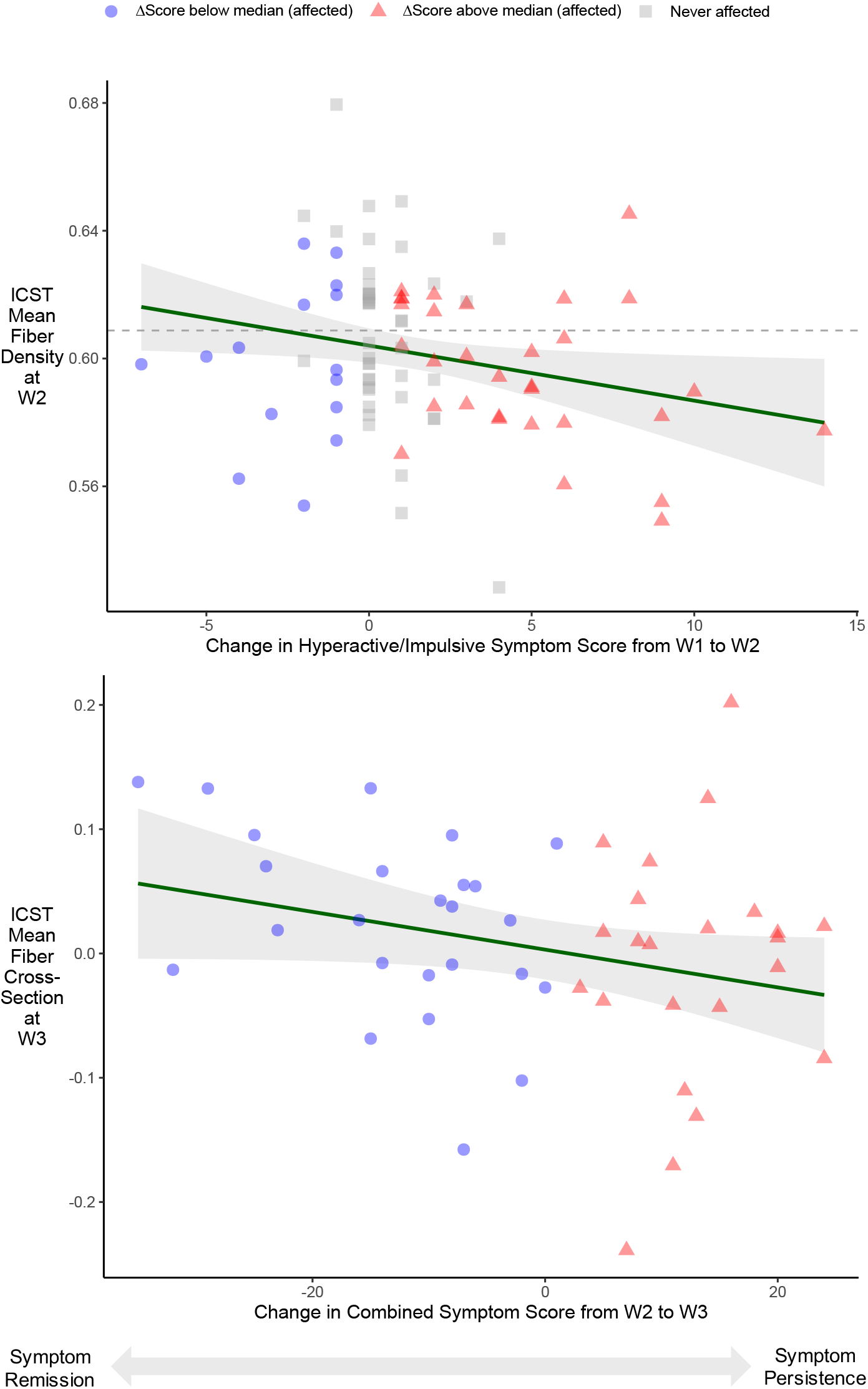
Graphs illustrating the relationship between left corticospinal tract (**Top**) mean fiber density values and change in hyperactive/impulsive symptom score from Wave 1 to Wave 2, and (**Bottom**) mean fiber cross-section values and change in combined symptom score from Wave 2 to Wave 3. Both plots include solid green regression lines with 95% confidence intervals. For reference, the mean fiber density at Wave 2 for unaffected participants is marked by a dashed gray line. Change in symptom score was calculated as: ΔScore = score _follow-up_ − score _baseline_. Participants are shown as individual points, differentiated according to whether their change in symptom score was above or below the median change in score.

**Figure S5.**
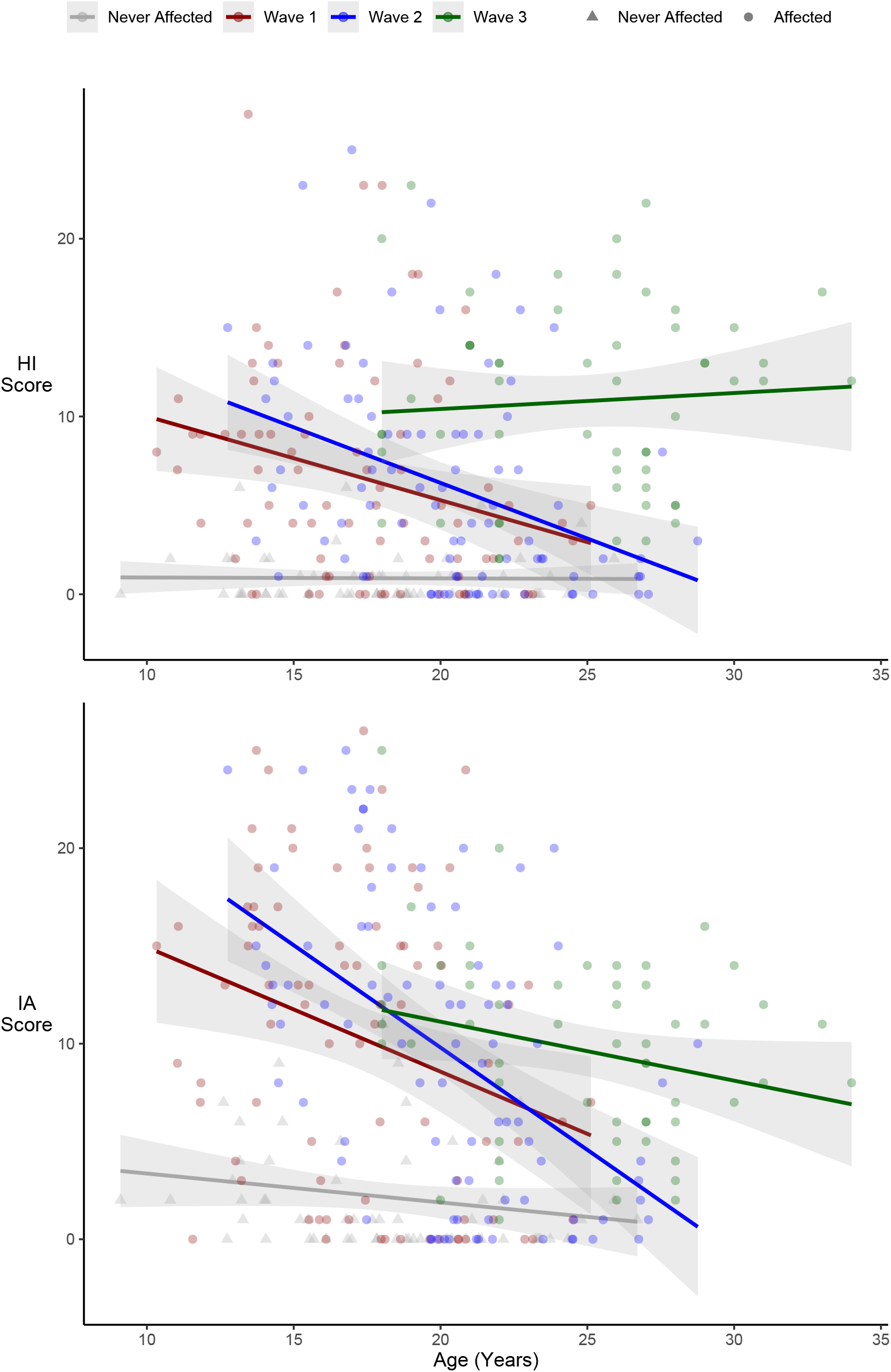
Participants from all waves pooled together (never affected are in gray triangles, and affected are circles colored according to wave in red [W1], blue [W2], and green [W3]) and plotted by age against symptom score in dimensions HI (**Top**) and IA (**Bottom**). Wave 3 included less participants and only those with a history of ADHD; scores were obtained with CAARS instead of CPRS.

